# Genomic and phenotypic characterization of an Oropouche virus strain implicated in the 2023-24 large-scale outbreak in Brazil

**DOI:** 10.1101/2024.08.02.606252

**Authors:** Elisa de Almeida Neves Azevedo, Alexandre Freitas da Silva, Verônica Gomes da Silva, Lais Ceschini Machado, Gustavo Barbosa de Lima, Bruno Issao Matos Ishigami, Keilla Maria Paz e Silva, Mayara Matias de Oliveira Marques da Costa, Diego Arruda Falcão, Andreza Pâmela Vasconcelos, Clintiano Curvêlo da Silva, Felipe Gomes Naveca, Matheus Filgueira Bezerra, Tulio de Lima Campos, Bartolomeu Acioli-Santos, Marcelo Henrique Santos Paiva, Clarice Neuenschwander Lins de Morais, Gabriel Luz Wallau

## Abstract

The *Orthobunyavirus oropoucheense* in a arthropod borne zoonotic pathogen known to infect sylvatic animals and humans by means of biting midges transmission. Several large-scale human outbreaks of Oropouche virus (OROV), primarily confined to the Amazon region, were documented over the decades. However, since 2022, more widespread OROV outbreaks have been unfolding in Brazil and across South America, with cases exported to Cuba, Italy, and Germany. I In Brazil, the virus has reached and established communitary transmission in states from all major regions of the country. Here we isolated, characterized the cytopathic effect and the full genomic sequence of two OROV isolates from the current outbreak. Our data shows that OROV can readily infect and replicate in non-human primate cells, supporting the role of non-human primates as important reservoirs. Phylogenetic data supports a direct introduction of the same lineages causing the 2022-24 outbreak in Brazil from the Amazonas state, the epicenter of the epidemics in Brazil. Lastly, as case counts accumulate in the state and the Northeast region, clear evidence supports established and sustained transmission. Continued studies are critical to understand the transmission cycle in this region, the most important vectors and reservoirs, to appropriately deploy control measures.

## Introduction

Oropouche Fever (OF) is a human disease caused by an arthropod-borne virus (arbovirus) of the genus *Orthobunyavirus*, family *Peribunyaviridae*, the *Orthobunyavirus oropoucheense* virus (OROV). This virus has a segmented, single strand negative sense RNA genome, making it susceptible to genetic reassortment. Despite its significant public health impact in South America, particularly in the Amazon region, OROV cases continue to be neglected and underreported^1^. Patients infected by OROV typically present non-specific symptoms common to arbovirus infections such as headache, myalgia, arthralgia, nausea, diarrhea. Hence case definition in regions with endemic circulation of other highly prevalent arboviruses (i.e Dengue, Chikungunya and Mayaro viruses) is only possible with the use of serological and/or molecular laboratory tests.

OROV was detected and isolated for the first time in Brazil in 1960, from a sloth (*Bradypus tridactylus*) blood sample ^2^ and since then, outbreaks have been documented across Amazonian countries ^2–4^. In Brazil, until the arrival of Chikungunya and Zika virus, OROV was the second more prevalent arbovirus infecting humans only after Dengue virus ^5^. Transmission of OROV occurs primarily through the bites of *Culicoides paraensis* ^6^ and potentially anthropophilic mosquitoes like *Culex quinquefasciatus* and *Aedes aegypti* ^7,8^. *However*, as *C. paraensis* also feeds on humans in urban settings during population bursts; it is likely the most important OROV vector. The broad distribution of these vectors in the Americas raises concerns of a high risk of widespread outbreaks.

Recent research has uncovered a new reassortant OROV strain circulating in Brazil since 2010-14, which has led to the ongoing outbreaks that began in 2022^9^. These outbreaks have expanded beyond the Amazon, affecting multiple Brazilian states ^10,11^. Of particular concern are the findings of OROV infection in multiple tissues from deceased fetuses in Pernambuco (https://www.gov.br/saude/pt-br/assuntos/saude-de-a-a-z/f/oropouche/notas-tecnicas,https://portalcievs.saude.pe.gov.br/noticias/DOCUMENTOS/arboviroses - reports Nº 21 and 22/2024), highlighting the potential for vertical transmission of this virus, potentially leading to adverse pregnancy outcomes. Moreover, recent findings identified the first adult deaths associated with OROV infection^12^. The present study reports the isolation of an OROV strain from 2023-24 outbreak in Brazil, including analysis of its cytopathic effect and phylogenetic relationships. These resources provide an important contribution to our understanding of this neglected arbovirus.

## Materials and Methods

### Samples and diagnostic

Serum samples from passive surveillance were collected in basic health units from patients showing arboviruses symptoms and forwarded to be tested at Laboratório Central de Saúde Pública de Pernambuco (LACEN-PE). Samples from patients up to 5 days after their first symptoms were processed for molecular tests. Total RNA extraction were conducted according to the manufacturer’s instructions, using Extracta® Kit-DNA e RNA de Patógenos MDx (Loccus, Cotia, SP, Brazil) for viral detection in clinical samples, or Maxwell 16 Viral Total Nucleic Acid Purification Kit (Promega Corporation, Madison, USA) for viral detection in cell culture supernatant. RT-qPCR was then performed using a kit multiplex manufactured by Instituto de Biologia Molecular do Paraná (IBMP) Lot 240984Z001 (IBMP, Curitiba, PR, Brazil). OROV, MAYV and MS2-specific primers and probes were employed, according to the manufacturer’s instructions. Reactions were performed with the ABI 7500 Real-Time PCR System (Thermo Fisher Scientific, Waltham, MA, USA) under the following cycling conditions: 45 °C for 30 min; 95 °C for 2 min; 94°C for 15 s; and 60 °C for 1 min (45 cycles). Clinical samples and culture supernatants were also evaluated for the existence of dengue, zika and chikungunya viruses, using Biomol ZDC II Kit (IBMP, Curitiba, PR, Brazil).

Ethical clearance for sample processing was approved by the Instituto Aggeu Magalhães ethical committee (CAAE - 10117119.6.0000.5190).

### Viral isolation from cell culture

Vero cells (ATCC® CCL81™) derived from kidney of an adult African green monkey (*Cercopithecus aethiops*) were cultured in Minimum Essential Medium (MEM) (Gibco/Invitrogen, CA, USA) supplemented with 10% (v/v) fetal bovine serum (FBS), 100 U/mL penicillin (Gibco/Invitrogen, CA, USA), and 0.1 g/mL streptomycin (Gibco/Invitrogen, CA, USA). Viral isolation was performed inoculating 50 μl of positive sera into Vero cells followed by maintaining the cells for 1 h at 37^°^C with 5% CO_2_. The cells were then further incubated for 4 days. Following this, OROV infection was confirmed by RT-qPCR.

### Virus growth and cytopathic effect

In a separate experiment, Vero cells were inoculated again in the two isolates obtained (hOROV/Brazil/PE-IAM4578/2024 and hOROV/Brazil/PE-IAM4637/2024) for the preparation of virus stocks which were collected upon detection of cytopathic effect and monitoring this effect. Supernatants from mock- and OROV-infected cellular cultivate were harvested at 24 hours post infection (hpi) and 48hpi and stored at −80 °C. Total RNA extraction and viral RNA detection was performed as previously described.

### Amplification and sequencing

Whole genome tailing amplification of OROV genomes and library preparation were performed using primers and the Illumina COVIDSeq kit according Naveca 2024 ^9^. Sequencing was performed using the MiSeq V2 reagent kit with 2×150 cycle paired-end run.

### Genome assembly and phylogenetic analysis

The genomes were assembled using a reference-guided strategy with the ViralFlow v.1.2.0 pipeline ^13^ for each of the three OROV segments (L - OL689334.1, M - OL689333.1 and S segments - OL689332.1). Sequences were deposited in GenBank (accession numbers: PQ073181-PQ073186). Phylogenetic analysis incorporated a dataset of complete OROV genomes, including new sequences from current/recent outbreaks (**Supplementary Table 1**). Alignments were conducted using MAFFT and trimmed with trimAl ^14^. A Maximum Likelihood phylogenetic tree was constructed using IQ-TREE, with the evolutionary model selected by ModelFinder and branch support was evaluated with SH-aLRT and ultrafast bootstrap with 1000 replicates. The generated tree was inspected using Figtree v1.4.4 (http://tree.bio.ed.ac.uk/software/figtree/).

## Results

### Virus isolation and cytopathic effect results

We obtained two OROV isolates from two patients with classic OROV-associated symptoms.

Analysis of the cytopathic effect of OROV on Vero cells was carried out by observing morphological changes in cells and cell death using optical microscopy. We observed that the cells began to show morphological changes from 24 hpi. At the 48 hpi time point, we observed destruction of the cell monolayer. Viral genetic material detection in the supernatant was confirmed as well as a decrease in Ct at 48hpi due to the increased detection of viral RNA.

### Genome assembly metrics, coverage breadth and depth

Both hOROV/Brazil/PE-IAM4578/2024 and hOROV/Brazil/PE-IAM4637/2024 isolates exhibited genomes with high coverage breadth (>95%) across all segments (L, M, S) (**Supplementary Table 2**). Furthermore the segments were showed a coverage depth between 6,330-10,003x (**Supplementary Table 2**).

Phylogenetic reconstruction revealed that the isolates obtained in this study belong to the same clade causing the large outbreak in the Amazon region (OROV_BR2015-2024_) reported by Naveca and collaborators 2024 ^9^. The sequences clustered with high node support to two sequences sampled in December 2023 from Manaus, the capital city of the Amazonas state (**Figure 2**). The two full genomes revealed 8 single nucleotide polymorphisms (SNP) between them (L=3, M=5 [**Supplementary Fig 1**]). Five SNPs are non-synonymous leading to amino acid changes, one for segment L (A5924T [M->L]) and four SNPs for segment M (C170T [T->I], G2530A [V->I], A2959G [I->V] and T3197C [V->A]). A comparison against the ancestral sequence of 2015 from Tefe, Amazonas (ILMD_TF29) repealed 73 SNPs for segment L, 45 for segment M and 11 for segment S, respectively (**Supplementary Fig. 2**).

**Figure 1.**
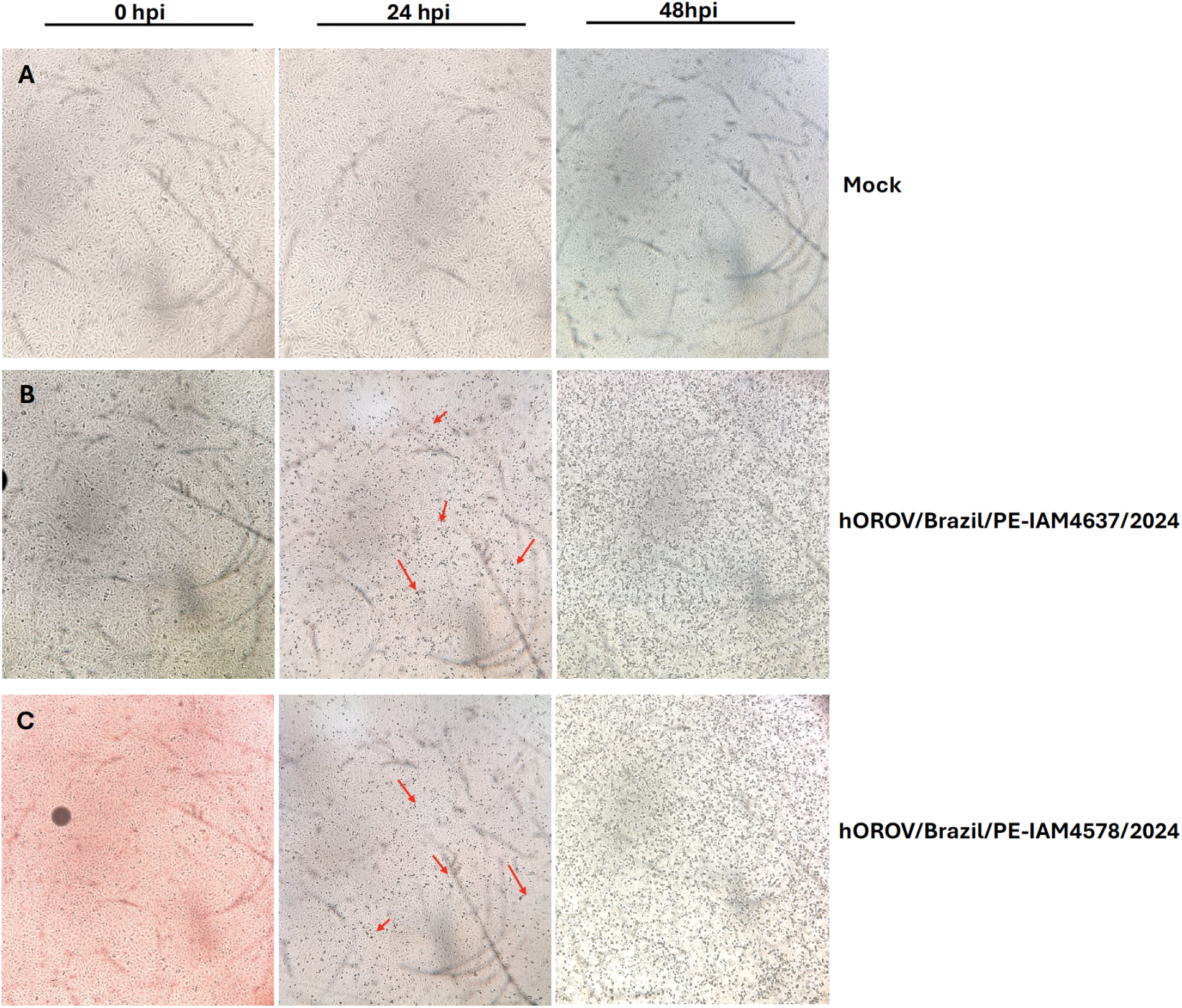
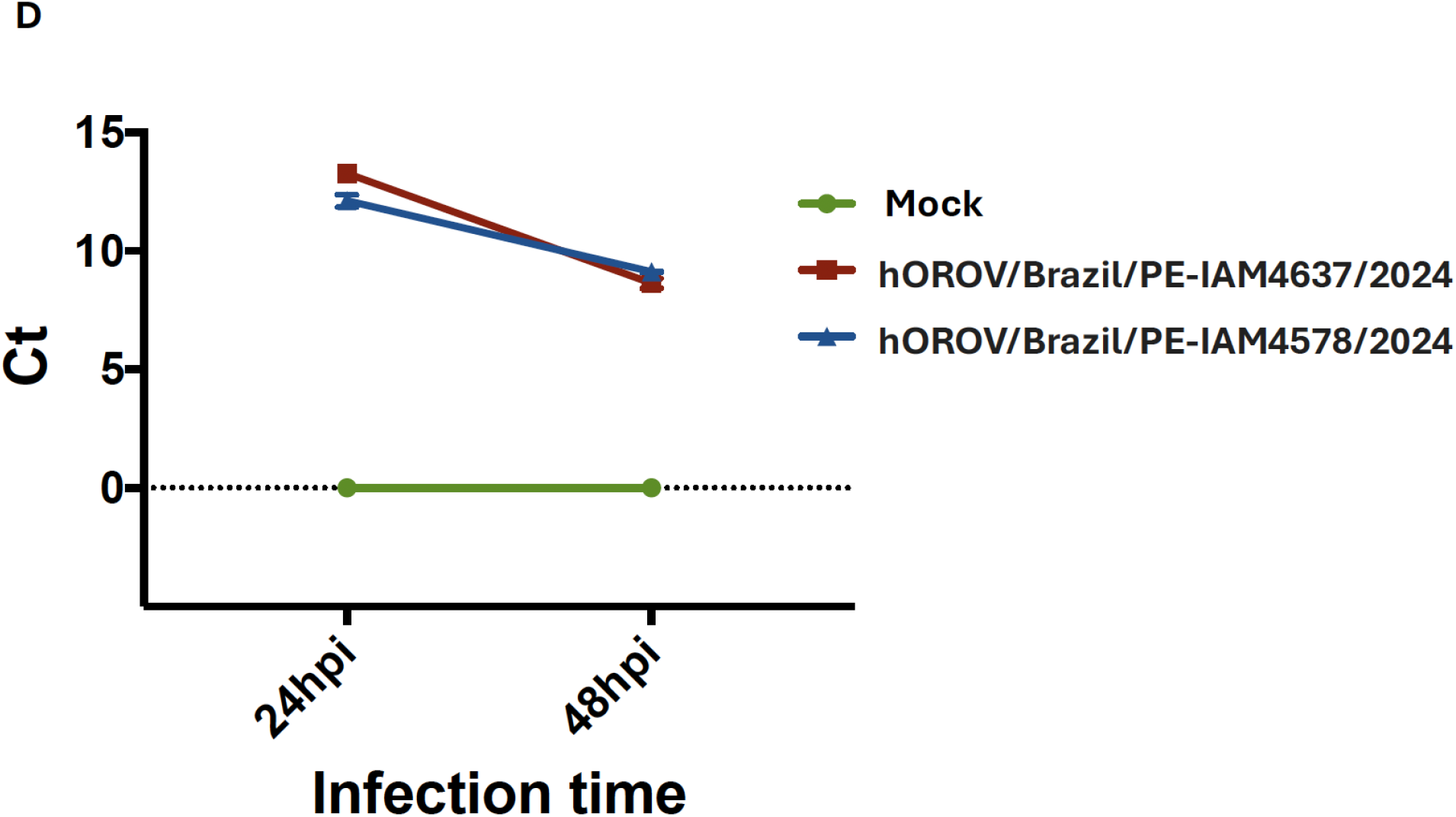
Light optical microscopy imaging of mock- and OROV-infected VERO cells at 48 hpi. Magnification 40X. (A) Uninfected cells used as infection control, were observed at the same times as infected cells. Left: first observed point, center: second observed point, right: third observed point. (B) Cells infected with OROV (hOROV/Brazil/PE-IAM4637/2024), were observed after infection (0 hpi on the left), 24hpi we observed the beginning of cytopathic effect (red arrows) and 48hpi we observed the viral cytopathic effect with destruction of cell monolayer (right). (C) Cells infected with OROV (hOROV/Brazil/PE-IAM4578/2024), were observed after infection (0 hpi on the left), 24hpi we observed the beginning of cytopathic effect (red arrows) and 48hpi we observed the viral cytopathic effect with destruction of cell monolayer (right). (D) Detection of viral RNA by RT-qPCR in the supernatants of cells infected with hOROV/Brazil/PE-IAM4637/2024 or hOROV/Brazil/PE-IAM4578/2024 and in control cells, at 24 hpi and 48 hpi. *hpi—hours post infection. Ct- cycle threshold*.

**Figure 2.**
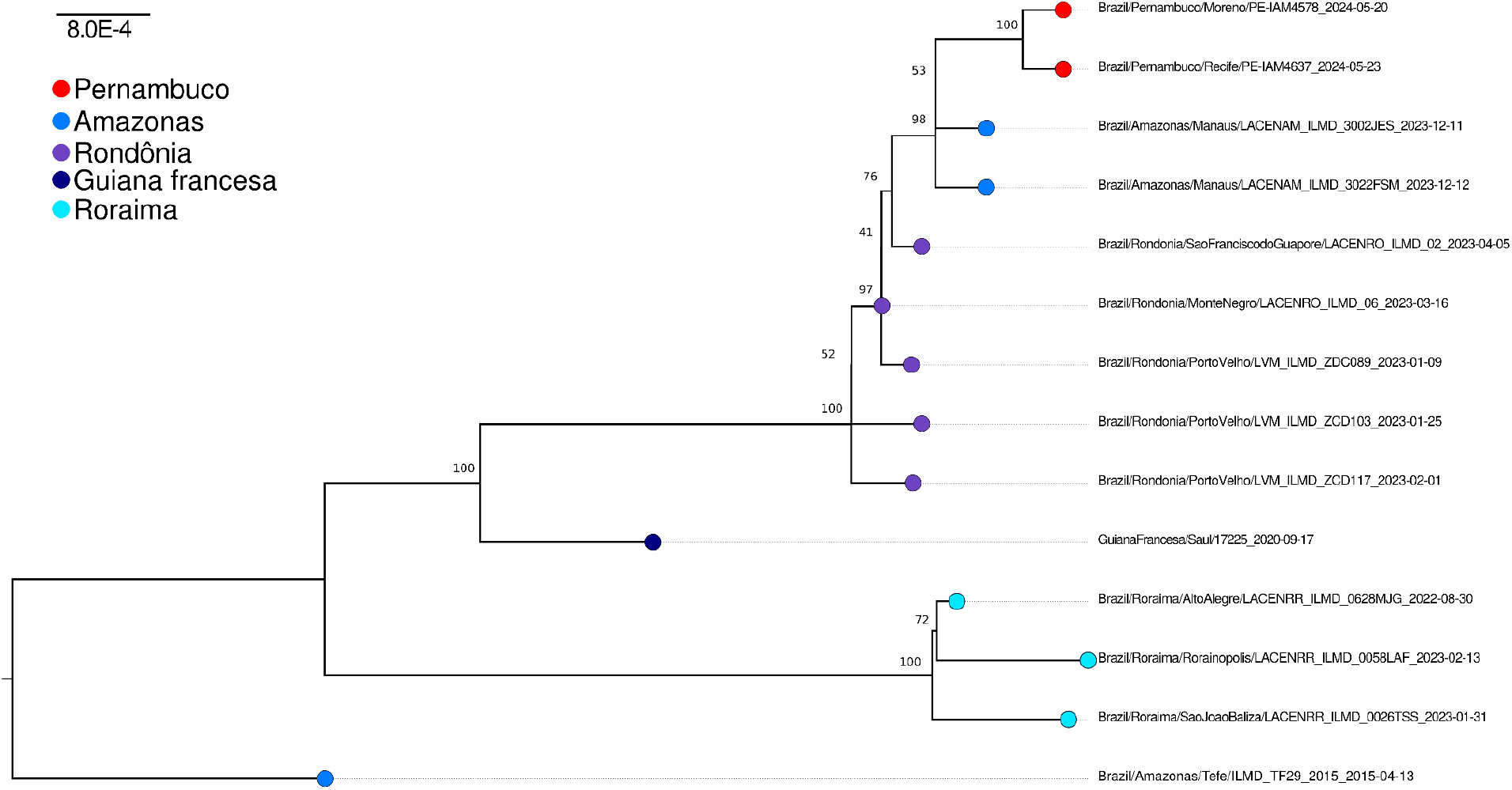
Maximum likelihood phylogenetic tree of two isolated Oropouche virus strains plus representative sequences from the current 2022-2024 outbreak. The phylogenetic tree was reconstructed based on a concatenated alignment of S, M and L segments of complete sequences available at the NCBI and analyzed on IQ-TREE 2.3.5 performing the SH-aLRT test and ultrafast bootstrap with 1000 replicates. Tip point colors represent the sampling location.

## Discussion

OROV is a neglected virus previously known to cause large human outbreaks in the Amazon region. Since 2022 there has been a new unfolding OROV outbreak that started in the Amazon region that now extended to all major Brazilian regions through exportation events and active communitary transmission. Here we isolated two strains in monkey kidney cells and characterized the genome of this strain. We found that the viral lineage isolated belongs to the same OROV_BR2015-2024_ lineage outbreak ^9^ from the Amazon region that emerged in Pernambuco from a direct importation event. *Culicoides paraensis* and *Culex quinquefasviatus* are an abundant vectors found in sylvatic-urban environments in Pernambuco. Both species as well other sylvatic mosquito species that were detected with OROV in the past (*Aedes serratus, Psorophora cingulata*, and *Haemagogus tropicalis*) ^4,15^ or closely related species circulating in the region such as *Aedes hastatus/oligopistus/serratus, Haemagogus janthinomys* and *Psorophora spp* ^16,17^ warrants further studies to reveal the main vector responsible for local transmission of the virus.

Scarce data exists about sylvatic reservoir species, but a number of widespread species of the South American fauna are known to be infected naturally by this virus including sloth, birds, rodents and non-human primates ^18–20^. Although we did not access sylvatic animals from active transmission areas in Pernambuco, sloths, rodents and non-human primates are abundant in the Atlantic forest of the Pernambuco state ^21,22^. Moreover, the relatively easy isolation of the virus in monkey kidney cells supports the hypothesis that non-human primates are susceptible reservoir hosts of OROV maintenance and amplification. This finding warrants further assessment of sylvatic animals’ natural infection in Pernambuco to understand if sylvatic animal transmission established in the state or human cases are occurring by a strictly human-to-vector transmission only. The replication kinetics of the isolated OROV strains in monkey kidney cells also supports the findings that viremia peaks very rapidly (less than 48 hours) in cell culture and human subjects highlighting that after infection, OROV replication is rampant leading to relatively short viremic periods (acute phase lasts for **2** to **7** days) ^23^.

This new isolated strain with associated full genome available will be a key resource to study and develop new diagnostic and pharmacological tools to reduce the impact of this obscure virus on the human population.

## Data availability

Wallau, Gabriel (2024). Genomic and phenotypic characterization of an Oropouche virus strain implicated in the 2023-24 large-scale outbreak in Brazil. figshare. Dataset. https://doi.org/10.6084/m9.figshare.26459578.v1

